# Molecular dynamics of spike variants in the locked conformation: RBD interfaces, fatty acid binding and furin cleavage sites

**DOI:** 10.1101/2022.05.06.490927

**Authors:** Deborah K. Shoemark, A. Sofia F. Oliveira, Andrew D. Davidson, Imre Berger, Christiane Schaffitzel, Adrian J. Mulholland

## Abstract

Since December 2019 the SARS-CoV-2 virus has infected billions of people around the world and caused millions of deaths. The ability for this RNA virus to mutate has produced variants that have been responsible for waves of infections across the globe. The spike protein on the surface of the SARS-CoV-2 virion is responsible for cell entry in the infection process. Here we have studied the spike proteins from the Original, Alpha (B.1.1.7), Delta (B1.617.2), Delta-plus (B1.617.2-AY1), Omicron BA.1 and Omicron BA.2 variants. Using models built from cryo-EM structures with linoleate bound (6BZ5.pdb) and the N-terminal domain from 7JJI.pdb, each is built from the first residue, with missing loops modelled and 45 disulphides per trimer. Each spike variant was modified from the same Original model framework to maximise comparability. Three replicate, 200 ns atomistic molecular dynamics simulations were performed for each case. (These data also provide the basis for further, non-equilibrium molecular dynamics simulations, published elsewhere.) The analysis of our equilibrium molecular dynamics reveals that sequence variation at the closed receptor binding domain interface particularly for Omicron BA.2 has implications for the avidity of the locked conformation, with potential effects on Omicron BA.1 and Delta-plus. Linoleate binding has a mildly stabilizing effect on furin cleavage site motions in the Original and Alpha variants, but has no effect in Delta, Delta-plus and slightly increases motions at this site for Omicron BA.1, but not BA.2, under these simulation conditions.

## Introduction

Since the SARs-CoV-2 pandemic began in the city of Wuhan in China at the end of 2019 the virus has spread around the world and caused, either directly or indirectly, an estimated 20 million excess deaths as of 31^st^ March 2022 (1). Between December 2019 and April 2022, SARS-CoV-2 has infected billions of people worldwide. This RNA virus has had ample opportunity to mutate during the course of countless replication events during this pandemic. The spike protein is one of the key viral proteins, being responsible for making first contact with host cells and indeed becoming the target of the host response. In this study we compare the structure and molecular dynamics of the spike protein of the variants beginning with that of the first sequenced SARS-CoV-2 (Original ‘Wuhan’ GenBank Accession MN908947.3), the Alpha variant first sequenced from Kent in England (lineage B.1.1.7), the Delta variant first sequenced in Africa (lineage B.1.617.2), the Delta-plus variant that caused widespread disease in India (lineage B1.617.2-AY1), and the currently dominant Omicron variants, focusing on the two parent Omicron lineages (lineages B.1.1.529-BA.1 ISL-6640916 and BA.2 QLD2568). We focus on the sometimes subtle effects of changes between these variants around the fatty acid binding site in the closed spike trimer, the interface between the closed receptor domains in the trimer and the furin cleavage site. We discuss how these changes may have contributed to altered spectra of symptoms and modulation of infectivity, particularly in the case of the omicron variants.

The SARS-CoV-2 variants have not elicited the same range of symptoms or disease severity as compared to the original virus from Wuhan. In the unvaccinated, there were progressive increases in both R numbers and disease severity from the Original to Alpha, Delta and Delta-plus (2) In the UK vaccinated individuals with residual resistance have found that the Omicron variants, though more infectious (3–5), have so far led to proportionally fewer hospitalizations and far fewer deaths. In the UK, which currently has around ~70% of the population vaccinated (2 vaccines, plus one booster) with Pfizer-BioNTech or Oxford-AZ vaccines, the ZOE COVID study (6) has revealed that symptoms, when present, are likely to be more cold-like than for previous variants. 82% of infected people experience runny nose, 70% fatigue, 69% sneezing, 69% headache, with only 53% of people reporting a persistent cough. On the 2^nd^ of April 2022, an estimated 4.5 million people had symptomatic COVID-19 in the UK (6). Countries that have largely avoided COVID outbreaks in the past, such as Hong Kong are now experiencing a massively surging omicron outbreak that is claiming lives. In Hong Kong in March 2022, the death rate per 100,000 population (3.44) exceeded that of the Delta outbreak in London in 2020 (3.17) and 67% COVID fatalities in the 5th wave in Hong Kong were unvaccinated (Hong Kong Centre for Health Protection and the Department of Health, GovUK).

### The spike protein

The ectodomain of the spike protein (see Figure 1) is a trimer, each monomer consisting of two domains, S1 and S2. The S1 domain is largely responsible for host interactions and contains a head region: an N-terminal domain (NTD) and a receptor binding domain (RBD) containing the ACE II receptor binding motif (RBM), which is also a major antibody binding epitope (7). Our team identified a free fatty acid (FA) binding pocket at the RBD interfaces of the locked SARS-CoV-2 spike that was occupied by the essential fatty acid, linoleate (LA^−^) in 2020 (8). Our previous simulations have shown that binding of LA^−^ in the FA site helps to stabilize a locked conformation (9) which appears more condensed than the LA-free closed form) and that the carboxylate head group of the FA makes consistent salt-bridge interactions with the R408, occasional interactions with K417 and persistent H-bonding interactions with Q409, across the locked subunit interfaces (10). Fatty acids have since been retrospectively discovered in the FA sites of spike proteins from SARS viruses that infect other species (11, 12). The S1 domain of the spike protein is responsible for attaching the SARS-CoV-2 virus to the host cell by means of interaction with host cell receptors. This is predominantly achieved through opening the conformation by raising one or more RBDs to expose the binding site for the human ACEII receptor (13). Additional host cell interactions have been identified, e.g the exposed, positively charged region resulting from furin cleavage at the S1/S2 junction (see below) binds to the human NRP1 receptor (14).

**Figure 1.**
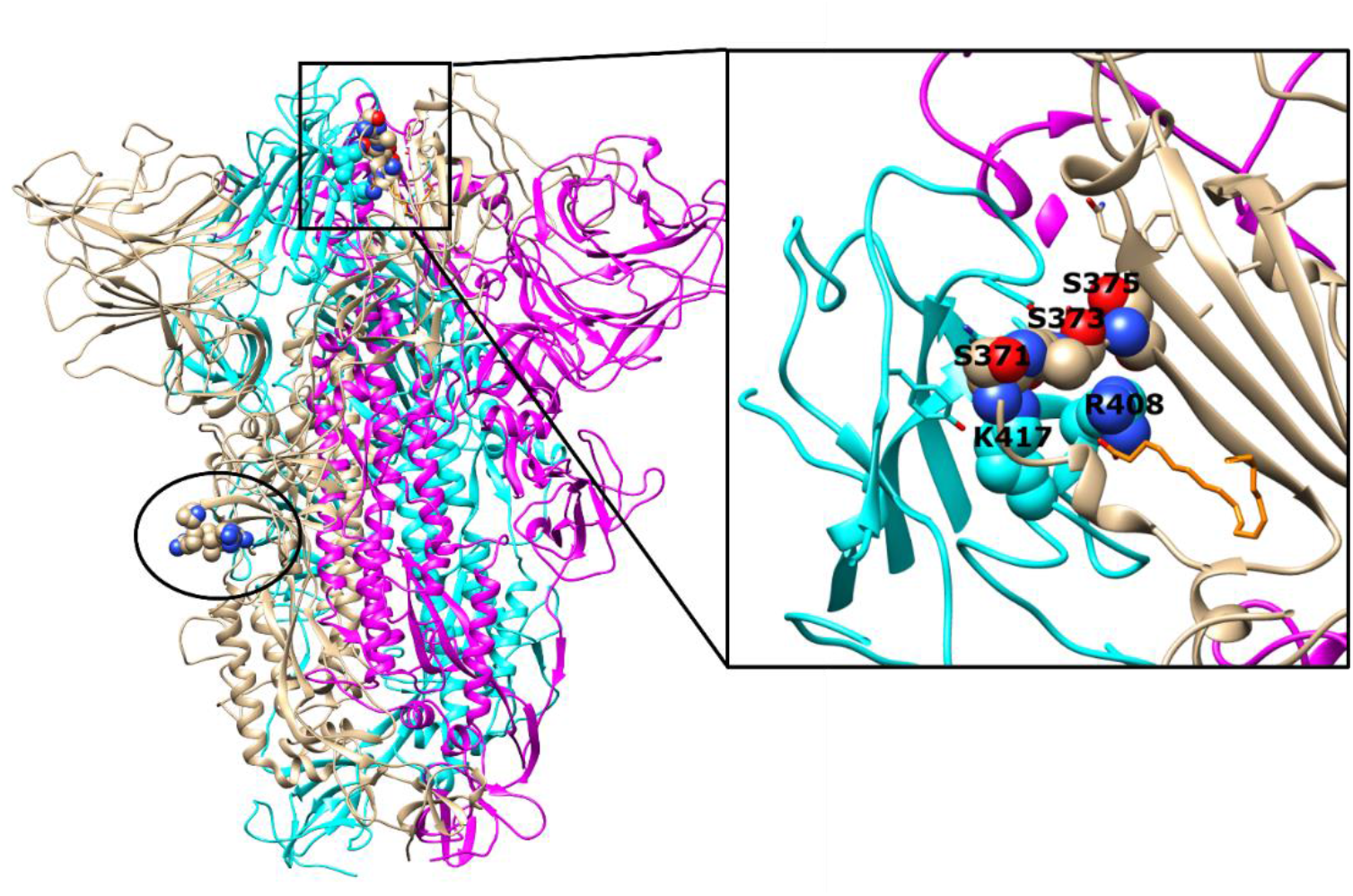
The Original (‘Wuhan’) uncleaved, locked, linoleate-bound spike trimer ectodomain showing the context for the furin cleavage site and changes at the RBD subunit interface occurring in the omicron variants. Glycosylation was removed for clarity. This structure is based on the cryoEM structures 6ZB5 and 7JJI with missing residues modelled from the first residue, and comprised of 45 disulphides per trimer, upon which all other variants in this paper are based. Spike chains are shown in cyan, wheat and magenta ribbon. The residues shown as spheres (circled) illustrate the position of the furin cleavage site. The inset panel is enlarged area showing residues as spheres where the RBD interface is changed in the omicron variants. Spike carbons are shown as spheres and coloured according to their chain, oxygens are in red and nitrogens in blue, linoleate (LA^−^) carbons are orange.

The S2 domain is required for membrane fusion events leading to cell invasion via the fusion peptide (FP), the fusion peptide proximal region (FPPR), two heptad repeats (HR1 and HR2) and the transmembrane domain (13, 15). The S1/S2 domains of the SARS-CoV-2 spike are joined by the furin cleavage site, which is unique among the betacoronaviruses. Cleavage by the human endoprotease furin, facilitates cell infectivity in humans and separates the two domains, revealing a positively charged region at the C-terminus of S1 (16). There is a further cleavage site in the S2 subunit (S2’) that is a target for the human TMPRSS2 protease and cleavage there allows membrane fusion to occur (16).

### Molecular dynamics simulations

All structures used here are based on the LA-bound, locked, unglycosylated, cryo -EM structure (6BZ5.pdb), using the N-terminal domain (NTD) from the NOVAVAX structure 7JJI.pdb, as more of the loop structure of the NTD was resolved in the latter. All the variants were modelled from this Original structure to maximise comparability of the molecular dynamics simulations. Missing residues of the NTD are modelled from the first residue, missing residues in loops have been modelled throughout and each trimer has 45 disulphide bonds formed. These models supercede the locked spike models we have used previously, which were missing the first 24 residues and had 42 disulphides (9, 10, 17). Model building and validation methods for models for this study are described in the supplementary material. These spike systems are relatively large (~600,000 atoms) and here equilibrium simulations were performed over 200 ns for three replicates for each variant. The authors note that extensive and noteworthy simulations have been carried out with glycosylated spike proteins (18–20) however, for comparing these variants, it was deemed that resolution of protein perturbations within the RBD and throughout the protein may be better achieved in the absence of surface-bound and highly flexible glycans, which may obscure the underlying protein dynamics over this short time-period. All glycosylation sites in the Original spike sequence as described by Zhao et al., (21) are retained in these variants. These equilibrium simulation data also provide the basis for dynamical non-equilibrium molecular dynamics simulations to map the allosteric pathways arising from the ligand annihilation from the fatty acid binding site in our accompanying paper (22).

All models were energy minimised and underwent short position-restrained molecular dynamics simulations in order to settle waters and generate three different sets of velocities for the three replicate production dynamics runs of 200 ns each, using GROMACS. Each spike trimer was simulated in a box of explicit waters with 150 mM NaCl, under periodic boundary conditions as an NPT ensemble at 310 K and pH 7, as described previously (17).

Sequence alignments are shown in Figure S1 and a phylogenetic tree in S2. Each Original and variant spike was subjected to three replicate simulations of 200 ns each in the presence of bound LA^−^s and with them removed (C-alpha RMSD plots for apo and LA^−^ bound are shown Figure S3). RMSF analyses over the trajectories of the apo and LA^−^ bound spike proteins (Figure S4) were carried in order to explore flexibility of certain regions of the variants in the presence and absence of LA, with particular focus on the RBD interface, the fatty acid binding site and the furin cleavage site.

**Figure 2** shows how the hydrophobic tail of LA^−^ lies in the hydrophobic pocket of one subunit in the Original locked spike trimer and the carboxylate head group interacts with charged (R408, K417) or polar (N409) residues on the neighbouring subunit, contributing to supporting a condensed closed (‘locked’) conformation.

**Figure 2.**
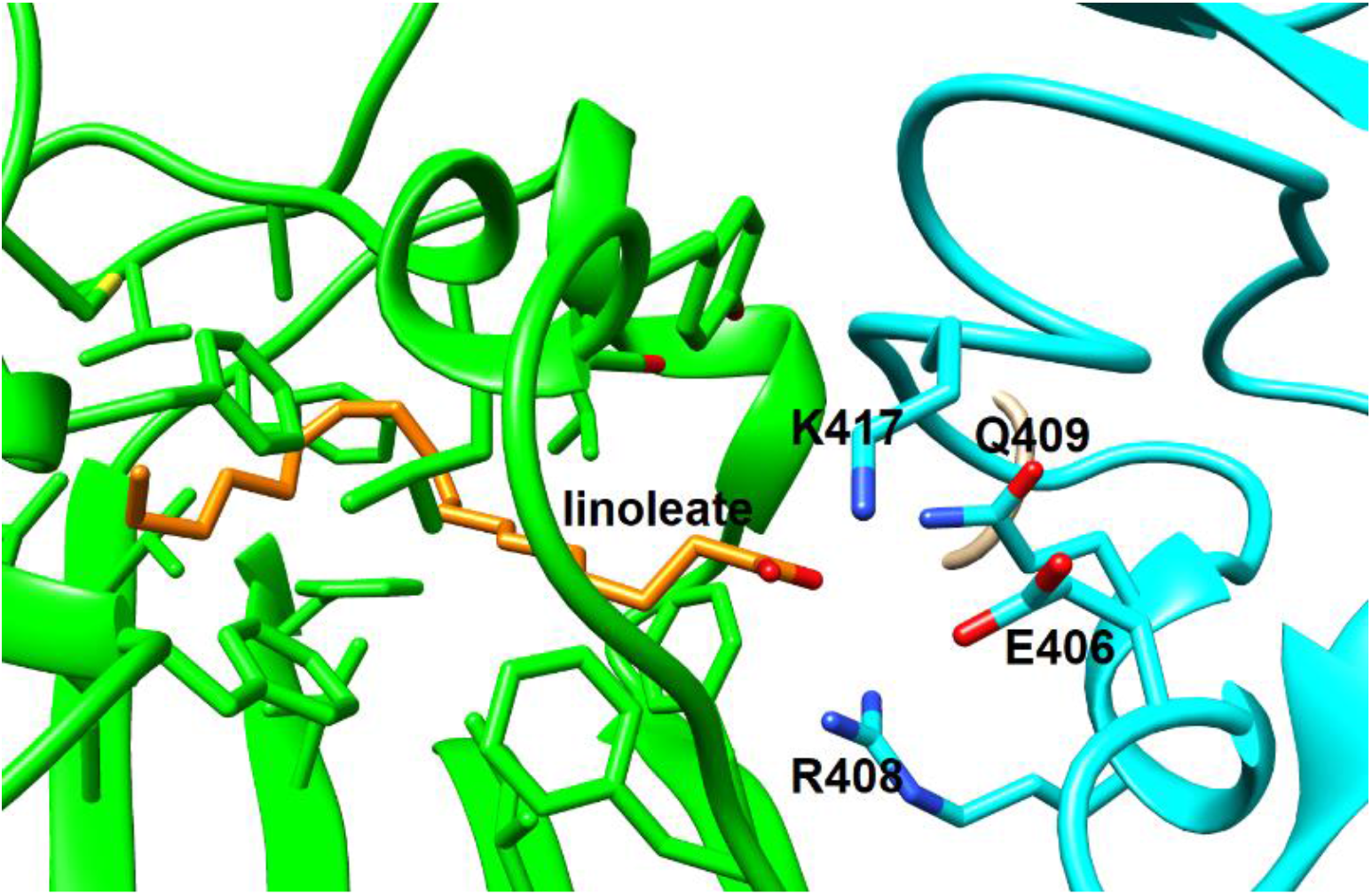
Close-up showing linoleate spanning a locked RBD subunit interface. This close-up shows the LA^−^ (depicted as stick, orange carbons) with its hydrophobic tail in the fatty acid pocket of RBD subunit A (green ribbon) and its carboxylate atoms interacting with residues (notably R408 and Q409) across the RBD interface on chain C (cyan ribbon). K417 and E406 are also shown for context. K417 makes occasional salt-bridging interactions with the LA^−^ carboxylates, but also interacts with E406 on the same subunit. Mutating K417 to asparagine (variant Delta-plus and Omicrons BA.1 and BA.2) reduces its ability to shield the repulsive negative charge of E406 (and D405) from the carboxylates of the LA^−^ whose hydrophobic tail is bound in the FA site of the opposite subunit.

#### Potential effects of sequence variation to the fatty acid binding site on linoleate binding

##### Original to Alpha

The residues lining FA binding pocket accommodating the hydrophobic tail of LA^−^ in B.1.1.7 (Alpha) remain unchanged from the Original. At the opposite end of the FA pocket, residues interacting with the fatty acid carboxylate also remain unchanged from the Original and maintain equivalent interactions to those of the Original R408, Q409 and K417 with adjacent residues Q414, T415, G416.

##### Original to Delta and Delta-plus

The residues lining the FA hydrophobic pocket in Delta are unchanged and residues interacting with the fatty acid carboxylate are also unchanged RQ (Original R408-Q409) and QTGK (Original Q414-K417). Likewise the fatty acid carboxylate interacting residues RQ (Original R408-Q409) remain unchanged in the B1.617.2-AY1 (Delta plus), however, the QTGK sequence is changed to QTGN, with the Original K417 changed to asparagine (N415 in Delta-plus). There is a loss of positive charge here, although the potential for hydrogen bonding is preserved, as discussed later. In the Original, K417 also provides occasional salt-bridge interactions with the carboxylates on LA^−^ and can shield them from the negative charge on the neighbouring E406 (E404 in Delta-plus), so there may be implications for the strength of the trans-subunit interactions, in the context of fatty acid binding, that seem to become progressively attenuated as the viral spike is mutated through Delta-plus, Omicron BA1 and Omicron BA.2, compared to the Original.

##### Original to Omicrons BA.1 and BA.2

Of all the major variants of concern to date, the Omicron lineages have diverged most from preceding ones with many more mutations throughout the spike protein (3). The FA pocket, in Omicrons BA.1 and BA.2, accommodating the hydrophobic tail of the FA is slightly modified by the substitution of (WT) S373 for proline. However, residues flanking this proline at the RBD subunit interface (see Figure 1) may affect the ease with which the RBD can be raised (9), as discussed later. In Omicron BA.1 the residues that interact with the LA^−^ carboxylate are equivalent to the Original residues R408 and Q409 (numbered R405 and Q406 in Omicron BA.1) but only Q406 is retained in BA.2. R408 is mutated to a serine in BA.2 (S405). The Original TGK located in positions 415-417 is TGN (T412-N414) for BA.1 and BA.2, as in the Delta-plus variant. The replacement of the positive charge at K417 by an asparagine retains hydrogen-bonding potential with the FA carboxylate and intersubunit H-bonding capacity, albeit reduced as discussed below. The substitution of the Original R408 for a serine in Omicron BA.2 (S405) has potential implications for LA^−^ binding affinity. R408 is the most dominant charged interaction with the carboxylate head group of LA in the FA binding site, across the RBD interface in the Original locked spike and this interaction persists for the nine LA molecules in the three replicate 200 ns simulations for 99.2% of the time. Interactions between all LA^−^s with their equivalent arginines persist for 98.6% in Alpha (R405), 99.9% in Delta (R405), 99.8% in Delta-plus (R406) and 99.7% in Omicron BA.1 (R405), of the simulation time

#### Exploring the effects of sequence changes to the locked RBD trimer interface in the Omicron variants for the LA^−^ bound (holo) simulations

Trans-subunit carboxylate interactions in the Omicron BA.2 simulations with LA^−^ bound (holo) were examined to establish whether other charged or polar interactions could compensate for the loss of the positive charge at R408 (S405 in BA.2). S405 fails to come within 5 Å of the LA^−^ carboxylates at any time in any of the omicron BA.2 holo simulations. Interestingly, BA.2 does seem to gain a repulsive trans-subunit interaction whereby a conserved glutamate at 403 (WT E406) on the opposite subunit comes within 5 Å of the LA^−^ carboxylates in BA.2. Unlike the Original, the negative charge of E403 (WT E406) is no longer shielded by the positive charge of K417, which is an asparagine in omicrons BA.1 and BA.2. The arginine at R400 in BA.2 (R403 WT) occasionally approaches the carboxylates to provide trans-subunit salt bridge interactions for 3 of the nine LA carboxylates for 36% of the (holo) simulations. The glutamine, Q406 (WT Q409) provides sustained (87% of the time) H-bonding interactions for 8 of the 9 LA carboxylates in BA.2 (68% for the ninth). N414 (WT K417) provides trans-subunit H-bonding for 7 of the nine carboxylates for 83-100% of the simulations, (the remaining two, for 30-40% of the time) but crucially the N414 side chain cannot salt bridge and cannot counteract the repulsive influence of the negative charge of E403 (WT E406) across the RBD subunit interface. The remaining two spend less time within range as the trimer interface is less tightly locked (see figures 3 and 4).

**Figure 3.**
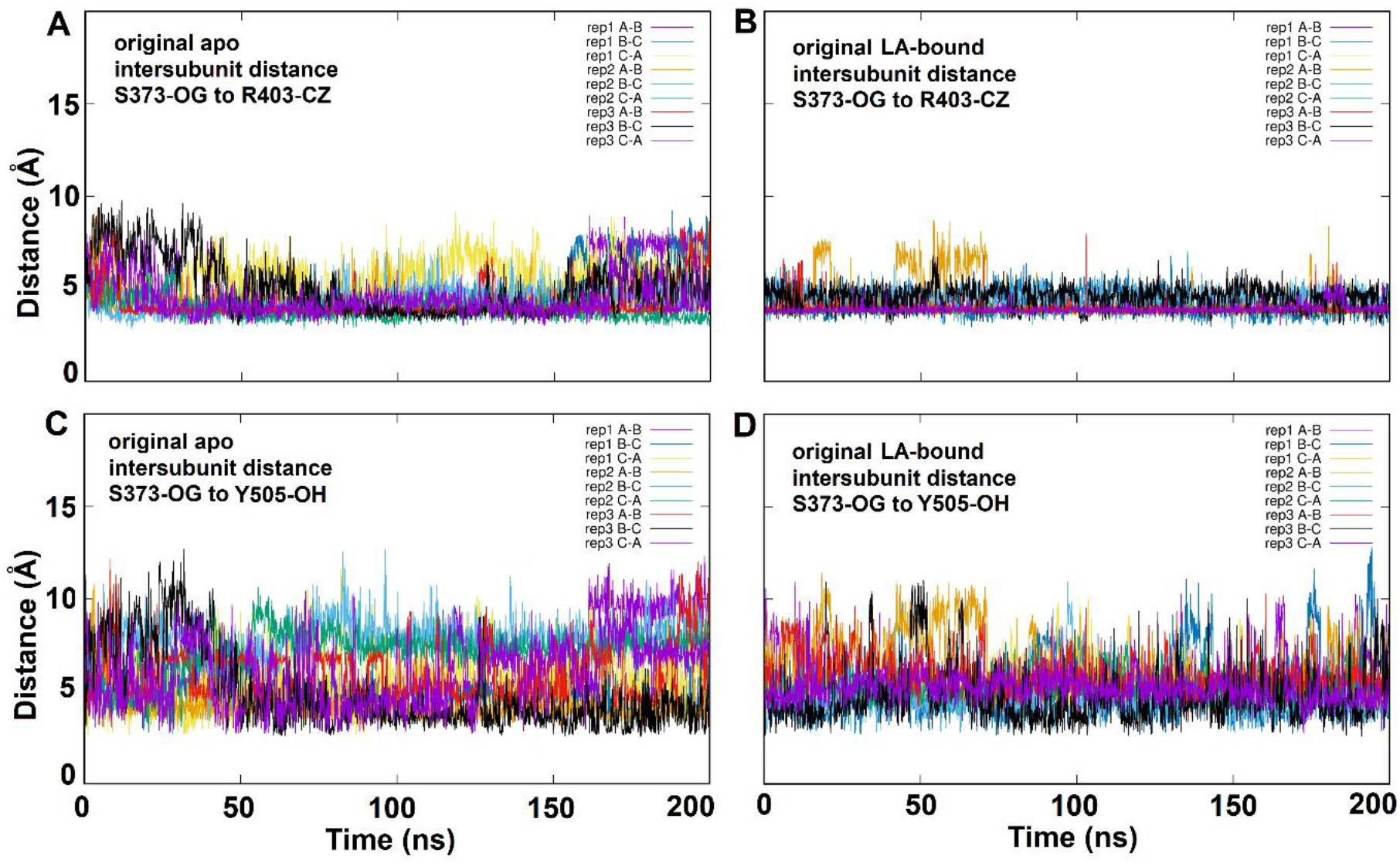
Interaction between S373 on one chain with R403 and Y505 of the neighbouring chain across the RBD interface in the Original spike trimer. Panel A shows the distances between the OG oxygen of S373 on one chain and the CZ carbon of R403 on the neighbouring chain for each interface of three 200 ns replicate simulations of the apo spike. Panel B shows the inter-subunit distances between S373-OG and R403-CZ when LA^−^ is bound. Panel C shows the distances between the OG oxygen of S373 of one chain and the OH oxygen of Y505 on the neighbouring chain for each interface of three 200 ns replicate simulations of the apo spike. Panel D shows the inter-subunit distance between S373-OG and Y505-OH when LA^−^ is bound.

**Figure 4.**
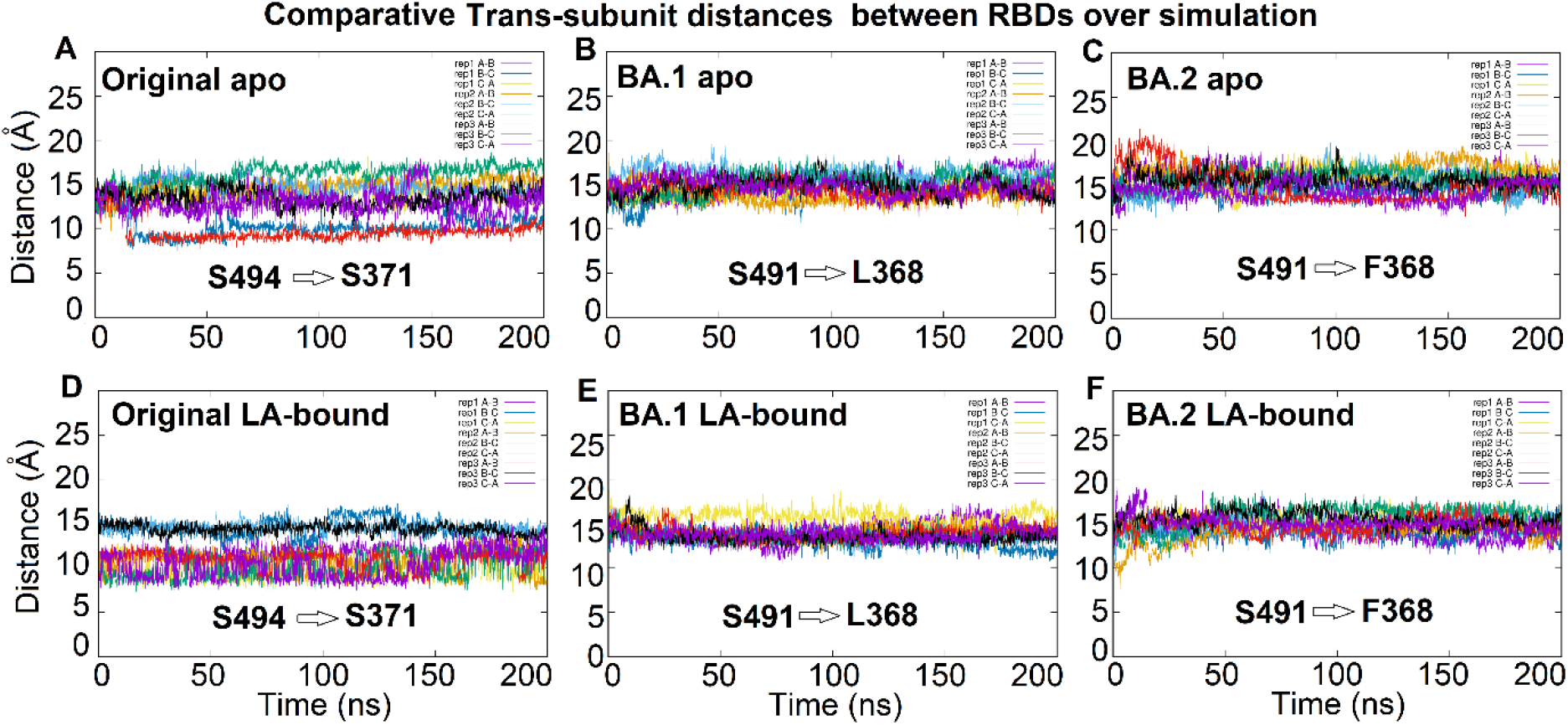
Trans subunit RBD distances during the MD simulations for the corresponding residues of the Original and Omicron variants in the presence and absence of LA. **A)** Inter-subunit distance between the Cα atom of S494 of one subunit with the Cα atom of S371 on the neighbouring chain for the Original apo SARS-CoV-2 spike during 3 replicate, 200 ns MD simulations. **B)** Inter-subunit distance between the Cα atoms of the mutated residues in Omicron BA. 1 (corresponding to the interactions between S494 and S371 of the Original), which in Omicron BA. 1 are between S491 and L368 (Omicron BA. 1 numbering) for 3 replicate, 200 ns simulations of apo spike. **C**) Inter-subunit distance between the Cα atoms of the mutated residues in Omicron BA.2 (corresponding to the interactions between S494 and S371 of the Original), which in Omicron BA.2 are between S491 and the mutated F368 (Omicron BA.2 numbering) for 3 replicate, 200 ns simulations of apo spike. **D)** Inter-subunit distance between the Cα atom of S494 of one subunit with the Cα atom of the S371 on the neighbouring chain for the Original LA-bound SARS-CoV-2 spike. **E)** Inter-subunit distance between the Cα atoms of the mutated residues S494 on one subunit and L368 in Omicron BA. 1 (corresponding to the interactions between S494 and S371 of the Original), for the LA-bound Omicron BA. 1 spike. **F)** Inter-subunit distance between the Cα atoms of the mutated residues S494 on one subunit and F368 in Omicron BA.2 (corresponding to the interactions between S494 and S371 of the Original), for the LA-bound Omicron BA.2 spike. Here we see that in the Original apo spike the trimer interfaces are a mixture of tightly closed and more relaxed states (A) and the presence of LA favours the tighter conformation (D), whereas for the Omicron variants all interfaces are more relaxed in the apo form (B&C) and the presence of LA makes no difference (E&F).

#### Effects of mutations at K417 (Original) to the RBD trans-subunit interactions

The residues in positions equivalent to the Original K417 are N415 (Delta-plus) and N414 (Omicrons BA.1 and BA.2). These simulations show that the H-bonding potential for the asparagine substitution for K417 progressively reduces the trans-subunit RBD H-bonding potential from the Original, to the Delta-plus and further still for both Omicron variants. The Original spike K417 side chain remains within inter-subunit H-bonding distance with all residues LYNSAS (WT 368-373) on the neighbouring subunit for between ~30 to 90% of the duration of all simulations in both the apo and LA-bound states. The analogous N415 (WT K417) in Delta-plus remains within H-bonding interaction distances with only the YN of the LYNSAS region in the neighbouring subunit and for just ~16-50% of the time in the apo and LA-bound simulations. This loss of inter-subunit H-bonding capacity is further reduced for the analogous N414 of omicron BA.1 which only occasionally contacts the tyrosine of this span, (now LYNLAP) for on average 16% of the time over all trajectories. As mentioned above, the short stretch of sequence LYNSASFS (residues 368 to 375) in the Original spike is varied only in the omicron variants to LYNLAPFF (BA.1) and LYNFAPFF (BA.2) (Figure S1). This stretch lies on the interface between two subunits required to release an RBD to be raised for binding to the ACE2 receptor (9, 23). In the Original spike, S373 (WT) interacts predominantly with R403, K417, E406 and Y505 across the locked RBD subunit interface. R403 stays within inter-subunit H-bonding distance of S373 for almost 100% of all Original apo and LA-bound simulations, with K417 and E406 maintaining contact for about 80%, Y505, about 50% of the time (see Figure 3).

Interaction distances between S373 on one RBD subunit and R403 and Y505 of the neighbouring subunits for the Original spike are shown in Figure 3. The substitution of these Original serines (S371, S373 and S375) by hydrophobic residues in the Omicron variants precludes hydrogen-bonding interactions from contributing to keeping the RBDs locked at this point. Indeed, under these simulation conditions, this loop can become deflected away from the interface by around 5 Å (Figure 4).

Figure 4 shows that the three RBDs appear less tightly packed in this region of the closed structure in the Omicron variants (panels B,C, E and F) compared to the Original spike (panels A and D). The distance between the Cα atoms of the intersubunit residues in the omicron variants (L368 on one chain – S491 on neighbouring chain) spend more time at the furthest extreme compared with the corresponding residues (S371 – S494) of the Original spike apo simulations. This slight perturbation seems more consistent in the presence of LA^−^ across this part of the RBD subunit interface for Omicron BA. 1. RBD subunit packing is similarly affected in omicron BA.2, although the presence of LA^−^ seems to make little additional difference in this region in this case.

#### Comparing linoleate mobility within the FA binding site between variants

Figure 5 compares the motion of individual linoleates within their FA pockets for the Original and variant spikes. The motion of LA^−^s in the Original spike show that whilst some remain in relatively fixed in position with all-atom RMSDs averaged to ~ 1 Å, three of nine exhibit greater flexibility, largely in their hydrophobic tail conformation. This behaviour is repeated in the B.1.1.7 variant where only one LA^−^ flexes its tail, but the all-atom averaged RMSD of the others is slightly greater than the more static conformations in the Original. The motion of linoleates in B1.617.2 and B1.617.2-AY1 follows a similar pattern. Linoleates in Omicron BA.1 appear to be slightly more mobile generally, but seem to exhibit less tail motion. The most striking difference is for Omicron BA.2 where all the LA^−^s are more mobile within the FA pocket. This is probably attrituble to the loss of the carboxylate-anchoring R408 (now S405). Despite this slightly slackened RBD subunit interface for the omicrons, none of the LA^−^s dissociate from the spike under these conditions and on the (short) timescale investigated here.

**Figure 5.**
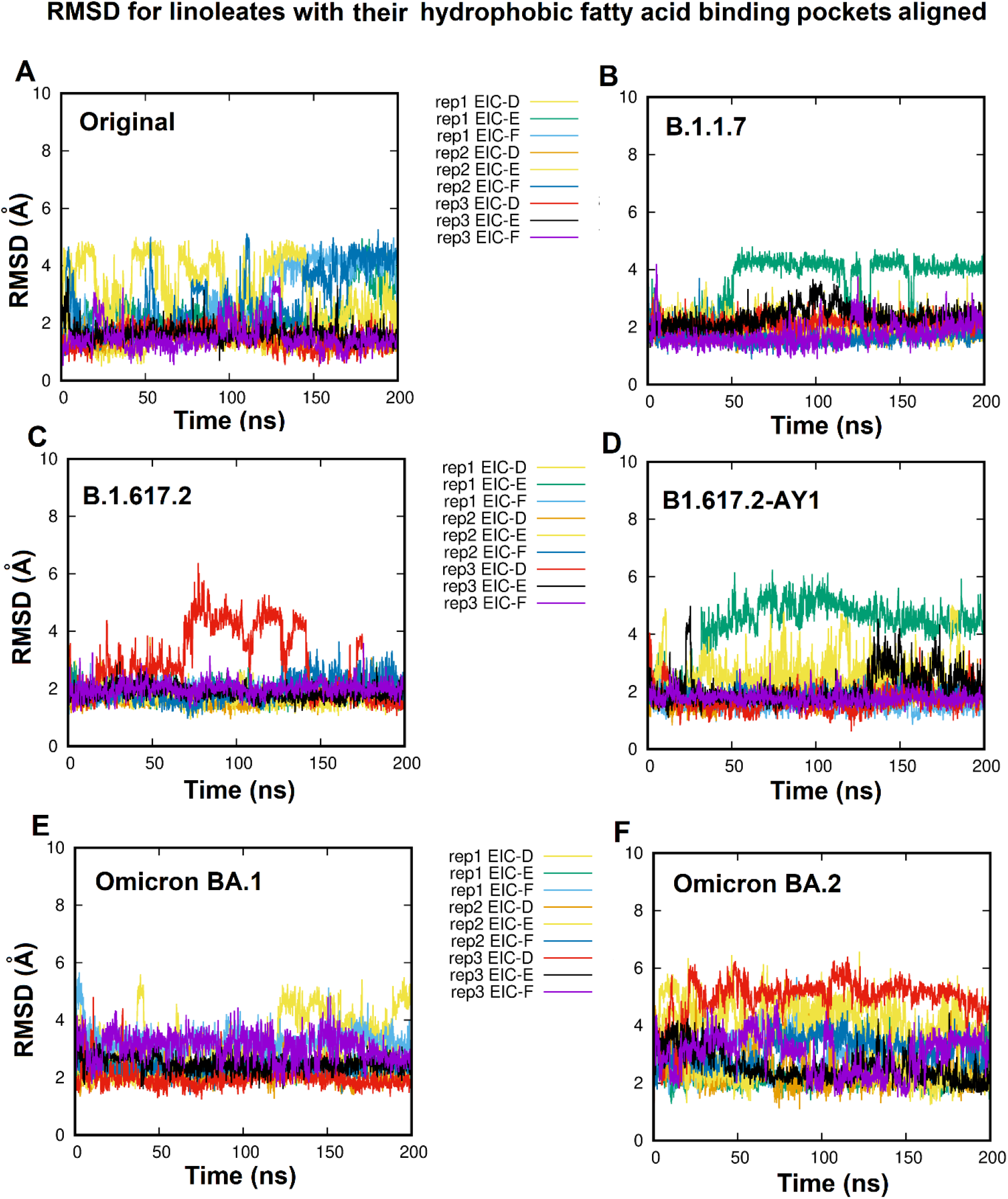
RMSD of individual linoleates for each replicate (200 ns) dynamics simulation, with the C-alpha atoms of their hydrophobic pockets aligned, to illustrate the motion of the linoleates within their pockets for (A) the Original, (B) the B.1.1.7 (Alpha), (C) B1.617.2 (Delta), (D) B1.617.2-AY1 (Delta-plus), (E) omicron BA.1 and (F) the omicron BA.2 variants. (The legend coloring is the same for all plots.) N.B. None of the LA^−^molecules leave the FA site during these simulations.

#### Effects of sequence variation on the furin cleavage site

The Original spike furin cleavage site spans residues PRRAR (Original P681-R685). The Original proline residue is mutated in all the variants discussed here. The furin active site has four negatively charged residues so it is likely that the addition of extra positive charge around the substrate cleavage site may increase binding affinity and/or cleavage rate by furin. The Original SARS-CoV-2 furin cleavage sequence was not ideal, but the addition of flanking positively charged residues may be making cleavage more efficient (24). In the alpha variant, the proline is mutated to a histidine, as is the case for Omicron. Histidine on an external loop may also become protonated at low pH. The loss of the proline alters O-glycosylation in this region which has been postulated to affect loop exposure to increase cleavage (25). For both the delta variants the potential for increased positive charge here was made permanent by the mutation of the Original proline for an arginine. Alpha, Delta and Delta-plus have been progressively more severe forms of COVID-19 and this may in part have been influenced by upregulated furin cleavage. This cannot be said for the Omicron variants. Clinical/epidemiological data show that infection by Omicron BA.1 leads less frequently to the severe disease outcomes elicited by Alpha or Delta variants in vaccinated populations. Though this may not always be the case in naïve populations. The analysis of the RMSF of the furin cleavage sites suggest that the furin cleavage sites of the apo forms of the variants are no more inherently flexible under these simulation conditions than the Original spike (Figure S5). T-tests performed to compare the RMSFs for each furin-cleavage site between their respective apo and LA^−^ bound forms can be found in table S1. Comparisons of furin-cleavage-site RMSF data for the Original apo with each of the variant apo forms can be found in Table S2. Finally furin-cleavage site RMSF data comparisons of the Original LA^−^ bound form with the LA^−^ bound forms of the variants can be found in Table S3. The caveat here is that the effects of glycans on these sites should also be considered. Our equilibrium simulations support our previous data (Original spike starting from residue 25) (10) suggesting that LA^−^ may have a weakly stabilising effect on fluctuations at the furin site in the full-length, Original spike. These data suggest this stabilising effect may also occur in the Alpha variant in the presence of LA^−^ in the FA site, but not for the Delta, Delta plus or Omicron variants. Indeed, for Omicron BA.1, bound LA^−^ may even increase fluctuations, which may influence the rate of furin cleavage. (Figure S7 and Tables S1 and S3).

## Discussion

The sequence variations of the spike proteins of different variants may affect their respective infectivity, and perhaps disease severity. The initial encounter with the host cell in the oro-nasal cavity may prove a crucial step in deciding the fate of the infection. The ease with which the spike RBD can open in the absence or presence of a fatty acid in the FA site is probably influenced by differences in directly interacting residues and the intersubunit interactions that hold the trimer closed. Our previous work (9) has shown that greater force is required to open an RBD in the Original spike when LA is bound. This was more pronounced for the BrisΔ (furin-cleavage site deletion). We surmise then that bound LA^−^ helps to reduce the frequency of spike RBD opening and consequentially, once bound and with the RBDs closed, LA is more likely to be trapped within and effectively sequestered. The ability of LA^−^ to limit RBD opening when occupying the fatty acid site would at first seem counter-productive to the virus. However if that fatty acid is LA^−^ sequestered from membranes of cells lining the oronasal cavity, which provide the first line of defence, this may benefit the virus by helping to subdue an initial, robust response as LA^−^ promotes an inflammatory response in the upper airways (26). It is perhaps worth noting that linoleic acid cannot be produced by the body, therefore availability within membranes relies on dietary intake and probably varies between individuals. The relatively small loss of hydrogen-bonding across the RBD subunit interfaces for the Omicron spikes (LYNLAP/LYNFAP for LYNSAS) in conjunction with N414 (Originally K417), may subtly ease RBD opening. We see from these equilibrium simulations that the closed RBD spike interface is less tightly packed in the Omicron variants than for the Original closed spike, suggesting that this subunit interface more readily dissociates to open. If the Omicron variants exhibit a higher frequency of RBD opening due to mutations at the closed RBD interface, then more frequent openings would lead to more frequent, productive, ACEII interactions, hence increased infectivity and the corollary is that being more often open, they may also be less efficient at trapping or sequestering LA from cell membranes of the oronasal cavity. A more sparing effect on LA^−^ would seem to be supported by the observation that Omicron infections elicit more cold-like symptoms in the host (27) than previous variants. This could allow for a more immediate, robust and better contained inflammatory response within the upper airways, especially in those with some pre-existing immunity due to infection by previous variants, vaccination, or both. In Omicron BA.2, LA^−^ sequestration may be further hampered by the substitution of the dominating carboxylate interacting residue R408 (WT) for a serine (S405). S405 fails to come within 5 Å of the LA^−^ carboxylates in any of the omicron BA.2 holo simulations and our simulations reveal enhanced LA^−^ motility within the FA binding site, compared to all the other variants in this study. Reinfections have been problematic recently (28). Serine at the equivalent position to 408 may also potentially compromise the binding of some neutralising antibodies that would otherwise interact with R408 (29) and may facilitate re-infection with BA.2 following infection with BA.1, which is deemed possible but rare, according to a recent Danish study (30). Perhaps in the case of Omicron BA.2 key mutations resulting in a reduced susceptibility to antibodies may also reduce the need to sequester LA^−^. A combination of reduced LA^−^ sequestration, slightly reduced RBD interface affinity to ease opening, coupled with an escape-enhancing mutation at (WT) 408 could contribute to the increased infectivity, altered symptom spectra and enhanced immune evasion seen with Omicron BA.2.

## Acknowledgements

DKS and AJM thank BrisSynBio, a BBSRC/EPSRC Synthetic Biology Research Centre (Grant Number:BB/L01386X/1) for funding. DKS, ASFO and AJM thank EPSRC and HECBioSim (hecbiosim.ac.uk) for providing ARCHER/ARCHER2 time through a COVID-19 rapid response call. MD simulations were carried out using the computational facilities of the Advanced Computing Research Centre, University of Bristol (http://www.bris.ac.uk/acrc) under an award for COVID-19 research and the Oracle Public Cloud Infrastructure (https://cloud.oracle.com/en_US/iaas) under an award for COVID-19 research. We also thank the [Bristol UNCOVER Group](https://www.bristol.ac.uk/research/impact/coronavirus/research-priorities/) and the University of Bristol, for their support. AJM and ASFO thank EPSRC (grant number EP/M022609/1), BBSRC (grant number BB/R016445/1) and ERC (grant PREDACTED https://cordis.europa.eu/project/id/101021207) for support C.S. and I.B. are Investigators of the Wellcome Trust (210701/Z/18/Z; 106115/Z/14/Z).

## Author contributions

Conceptualisation/design of the work: DKS, ASFO and AJM; Acquisition, analysis and interpretation of the data: DKS; Writing of the manuscript: DKS, ASFO and AJM. Reviewing the manuscript: DKS, ASFO, ADD, IB, CS and AJM.

## Data availability statement

The data is openly available from MolSSI/BioExcel COVID-19 public data repository for biomolecular simulations (https://covid.molssi.org/simulations/).

## Notes

### Competing Interest Statement

The authors have declared no competing interest.

