## Supplementary_Material for "Molecular dynamics of spike variants in the locked conformation: RBD interfaces, fatty acid binding and furin cleavage sites"

#### Materials and methods

##### Equilibrium simulations

The sequence for the original (early 2020) spike was taken from Uniprot P0DTC2 (https://www.uniprot.org/uniprot/P0DTC2). The sequences for the alpha B.1.1.1.7, delta B.1.617.2 and delta-plus B1.617.2-AY1 were taken from UK-COG (https://www.cogconsortium.uk/). The sequences for omicron BA.1 and omicron BA.2 were taken from https://www.gisaid.org as EPI-ISL-6640916 and QLD2568 respectively.

The first 13 residues were modelled as extended stretch of peptide anchored at the first disulphide between residues C15 and C136 (original numbering) of the N-terminal domain (NTD) of the NOVAVAX structure (pdb code: 7JJI)(*1*), which had better defined loops for the N-terminal domain than previous structures (*2*). As such, the NOVAVAX NTD replaced the NTDs of our previous model (*3, 4*), which were originally based upon he cryo-EM structure 6ZB5 (*2*). For this study the original and variant models used for these simulations span residues (equivalent to the original protein) from 1-1139 and have 15 disulphides formed per monomer. Missing loops for the NTD modelled on the structure 7JJI (*1*), and the rest of the missing loops and deletions were modelled by hand. Each spike trimer had a total of 45 disulphides formed.

All variant models were adapted from the original protein (Figures S1-S2), sequence-checked with Clustal Omega (*5*) and checked using PROCHECK (*6*) for integrity and to ensure there were no cis-peptide bonds, or d-amino acids once loop modifications had been made. All models were considered acceptable if more than 99% of residues fell within allowed regions of Ramachandran space.

All models were energy minimised and underwent short position-restrained molecular dynamics simulations in order to settle waters and generate 3 different sets of velocities for the 3 replicate production dynamics runs of 200 ns each, using GROMACS (*7*). Each spike trimer was simulated in a box of explicit waters with 150 mM NaCl, under periodic boundary conditions as an NPT ensemble at 310 K and pH 7, as described previously in (*4*).

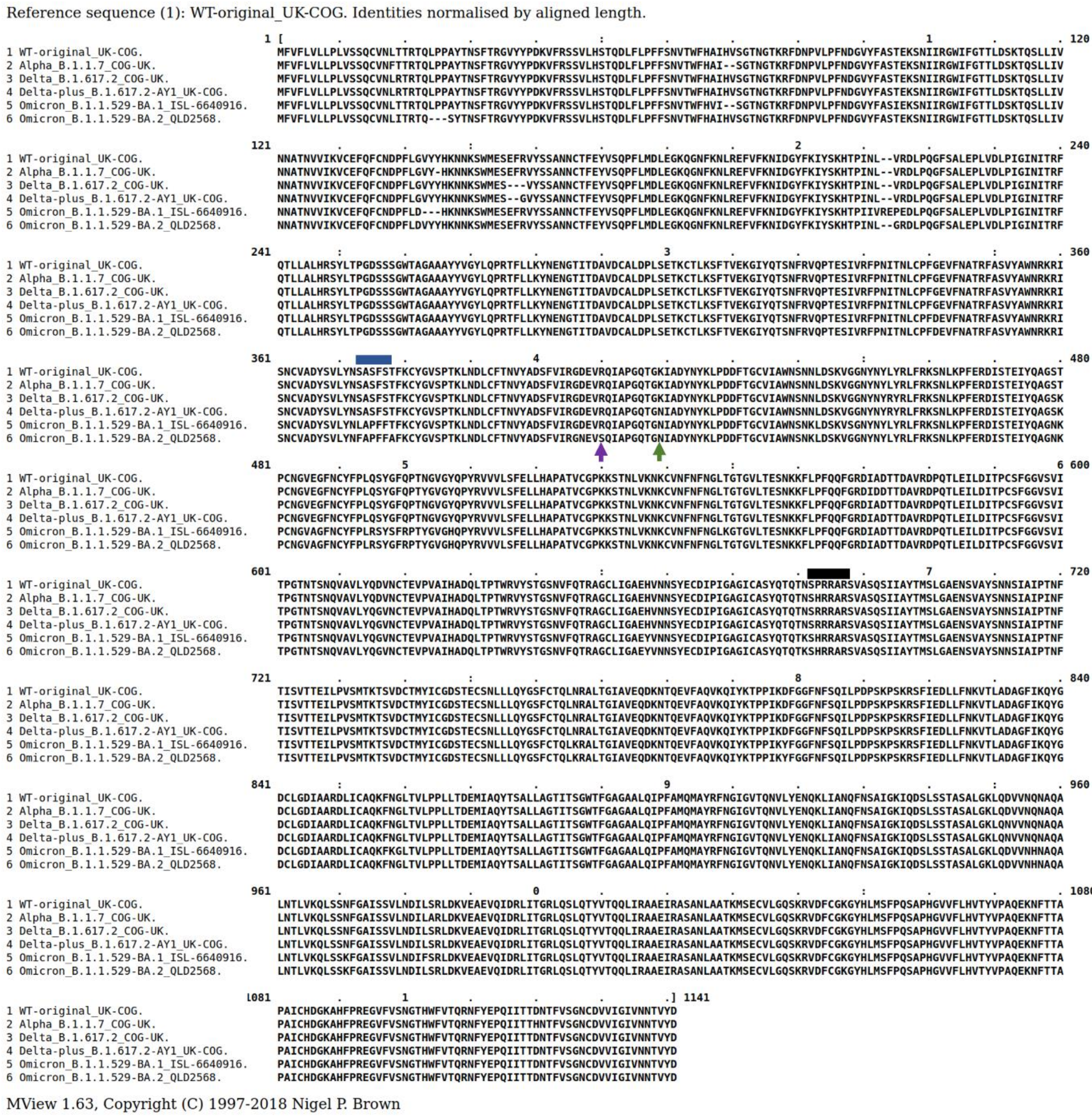


**Figure S1.** Shows a Clustal Omega sequence alignment (displayed by Mview 5 ) of the original SARS-CoV-2 spike protein with those of the alpha, delta, delta plus and omicron variants. The blue line indicates the mutations at the RBD interface in the omicron variants BA.1 and BA.2 that may affect the ease of RBD opening. The purple arrow indicates the position of the R408 in the original sequence that interacts with the carboxylate of LA. The green arrow indicates the position of the K417 in the original which interacts at the RBD subunit interface and the carboxylate of LA. The black line indicates the position of residues at the furin cleavage site in the variants.

Disulphides formed in all monomers (Original numbering):

C15-C136; C131-C166; C291-C301; C336-C361; C379-C432; C391-C525; C480-C488; C538-C590; C617-C649; C662-C671; C738-C760; C743-C749; C840-C851; C1032-C1043; C1082-C1126.

For information only, glycans not represented on these models.

Position of glycosylation sites (Original numbering):

N17; N61; N74; N122; N149; N165; N234; N282; T323/S325; N331; N343; N603; N616; N657; T678; N709; N717; N801; N1074; N1098; N1134 (*8*)

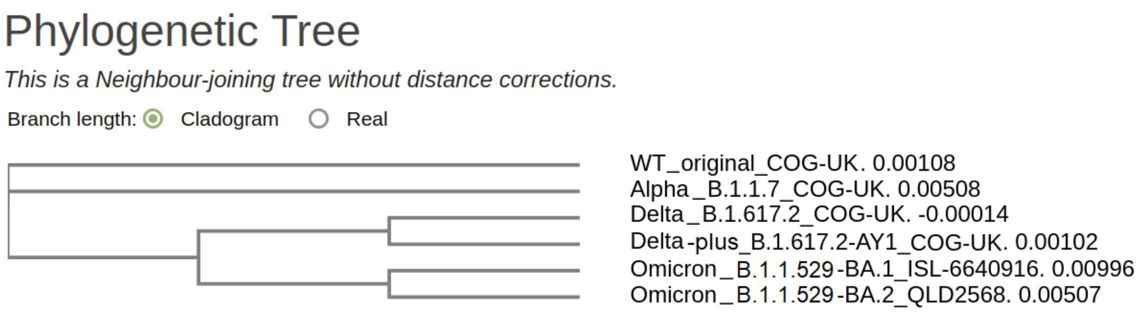


**Figure S2.** The phylogenetic relationship between the original and the alpha, delta, delta plus and omicron SARS-CoV-2 spike proteins (Clustal Omega (5)).

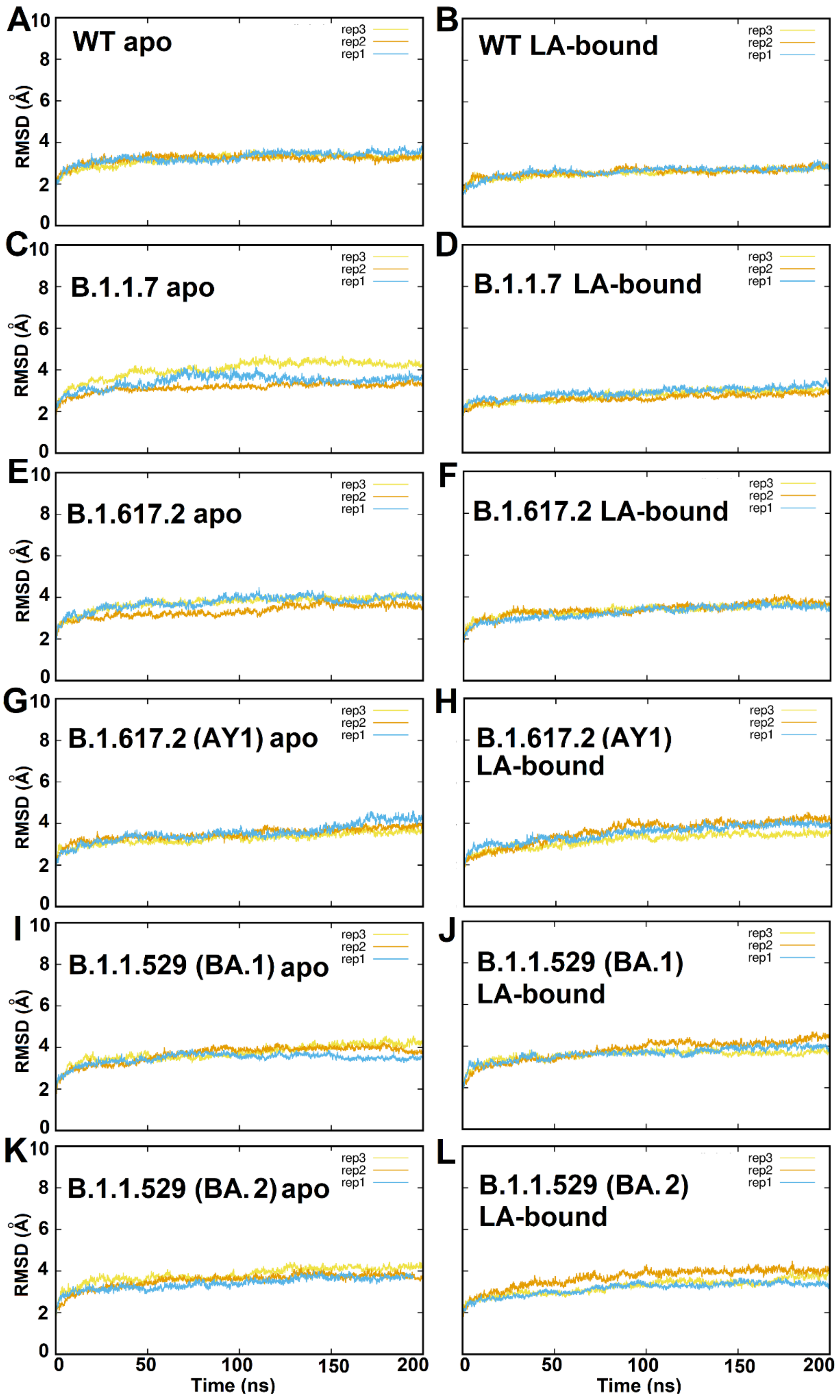


**Figure S3.** C-alpha RMSD plots for the equilibrium simulations compared to their starting structures.

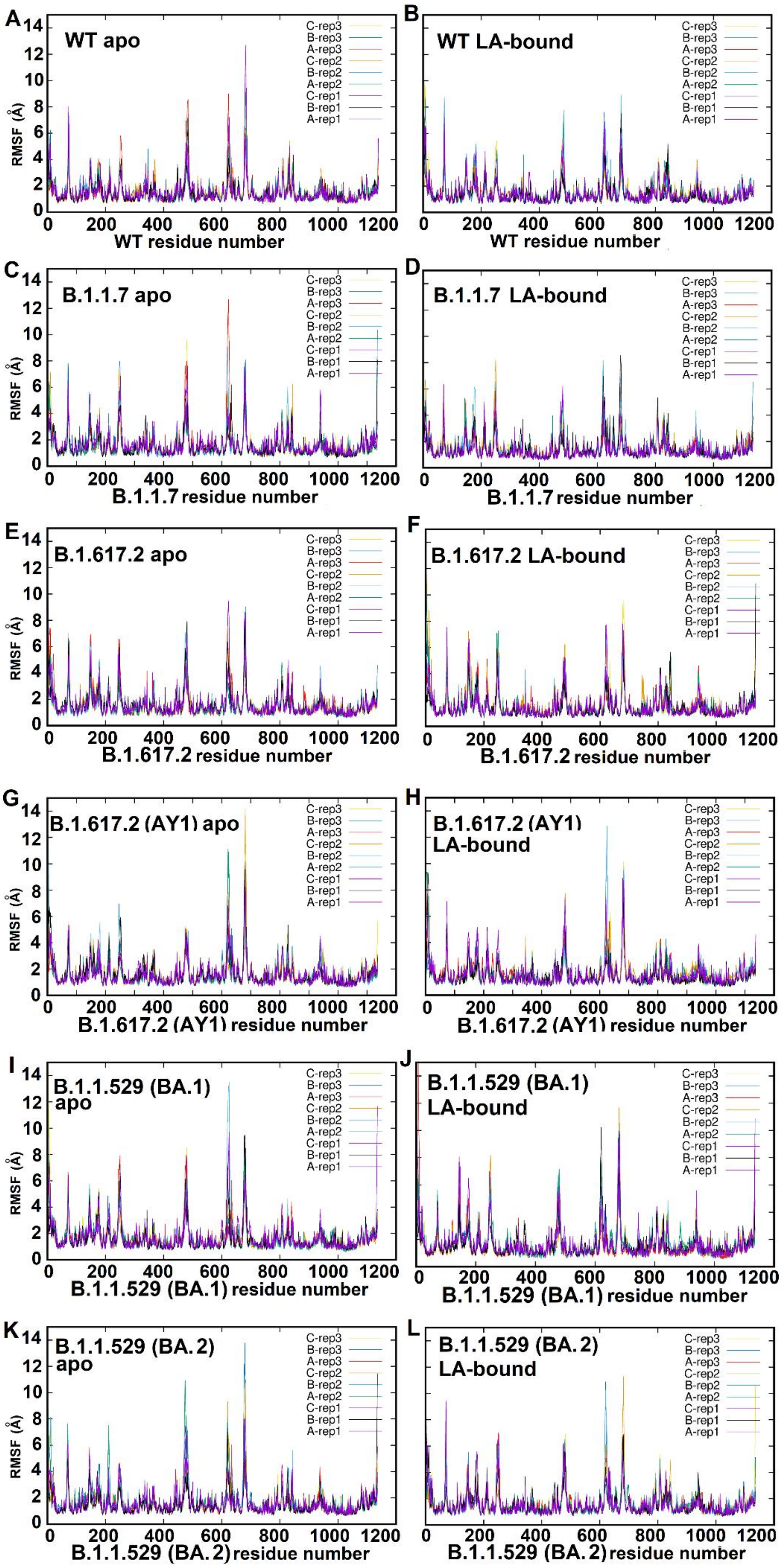


**Figure S4. Protein RMSF data for the original and variant spikes in the presence and absence of $LA^-$.** A) original apo spike. B) Original $LA^-$ bound spike. C) B.1.1.7 apo D) B.1.1.7 $LA^-$ bound E) B.1.167-2 apo F) B.1.167-2 $LA^-$ bound G) B.1.167-2-AY1 apo H) B.1.167-2-AY1 $LA^-$ bound I) omicron BA.1 apo J) omicron BA.1 $LA^-$ bound K) omicron BA.2 apo and L) omicron BA.2 $LA^-$ bound.

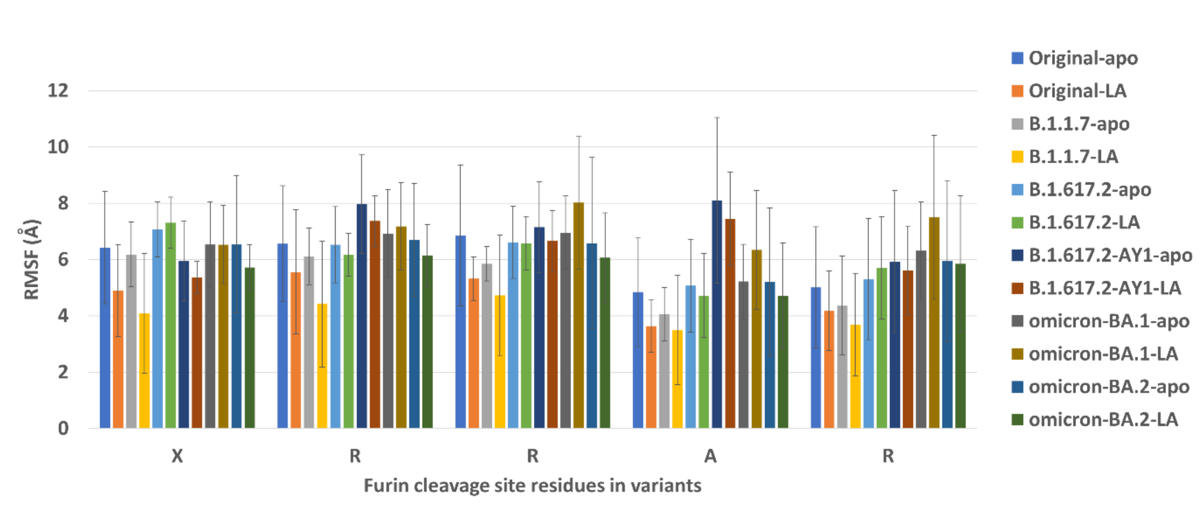


**Figure S5. The RMSF of furin cleavage site residues according to spike variant in the presence or absence of LA$^-$.** Fluctuations in this region appear to be stabilised somewhat by the presence of LA$^-$ in the original, and to a lesser extent in the B.1.1.7. LA$^-$ appears to have little effect on the furin site of B.1.617.2 and fluctuations which appear increased compared to the original. Fluctuations for -RRA- of B.1.617.2-AY1 are increased in the apo compared to the original and LA$^-$ has not reduced fluctuations back to that found in the original. For omicron BA.1, fluctuations in this region have not been reduced by LA$^-$, only increased. For omicron BA.2 fluctuations in this region, LA$^-$ appears to have had a slightly stabilising effect but remain higher than for the original or B.1.1.7 LA$^-$ bound simulations. The larger error bars are also inherently indicative of increased flexibility over the nine chains.

### Supplementary Tables

#### The effects of sequence variation on RMSF of furin cleavage site residues in the presence and absence of LA.

**Table S1. Paired (2-tail) t-tests comparing RMSFs for each variant's apo with its own LA$^-$ bound furin cleavage site.** Paired (2-tail) t-tests comparing the (protein) RMSF datasets for the furin cleavage site residues between the simulations with and without LA$^-$ bound for the original, alpha, delta, delta plus and omicrons BA.1 and BA.2 spike proteins. Each chain was treated separately so there were nine values for each of the residues over the 3 replicate 200 ns MD simulations for each apo variant and each LA-bound variant. Note that there is a weak significance (> 80% confidence) for three of the four residues' RMSF differences between the original apo spike with its LA$^-$ bound counterpart for residues P-RA-. The stabilising effect when LA$^-$ was bound was more significant for residues HRR—of the alpha variant but not for the last two residues ---AR. There was no appreciable stabilising effect of LA$^-$ on the furin site for delta, and not a significant effect on delta plus. LA$^-$ did not stabilise the furin site on omicron BA.1 but had a weakly significant (>74% CI) increasing effect on fluctuations of the –RAR. LA$^-$ appears to have had little effect on RMSF for the furin cleavage site in BA.2.

| Residue | Original-apo with Original-LA | Alpha-apo with Alpha-LA | Delta-apo with Delta-LA | Delta plus-apo with Delta plus-LA | Omicron-BA.1-apo with Omicron-BA.2-LA | Omicron-BA.2-apo with Omicron-BA.2-LA |
|---|---|---|---|---|---|---|
| **X** | 0.183 | 0.037 | 0.680 | 0.240 | 0.995 | 0.389 |
| **R** | 0.428 | 0.079 | 0.596 | 0.312 | 0.752 | 0.521 |
| **R** | 0.159 | 0.157 | 0.970 | 0.465 | 0.234 | 0.666 |
| **A** | 0.172 | 0.831 | 0.715 | 0.579 | 0.183 | 0.611 |
| **R** | 0.414 | 0.434 | 0.693 | 0.754 | 0.261 | 0.919 |

**Table S2. Paired (2-tail) t-tests comparing RMSFs for original apo with variant apo spikes' furin cleavage sites.** Paired (2-tail) t-tests comparing the (protein) RMSF datasets for the furin site residues between different variants without $LA^-$ bound. Note that the lowest values indicate the greater significance for the differences between the two datasets compared. Each chain was treated separately so there were nine values for each of the residues over the 3 replicate 200 ns MD simulations for each apo variant.

| | Original-apo with Alpha-apo | Original-apo with Delta-apo | Original-apo with Delta plus-apo | Original-apo with Omicron-BA.1-apo | Original-apo with Omicron-BA.2-apo |
|---|---|---|---|---|---|
| **X** | 0.777 | 0.388 | 0.599 | 0.905 | 0.915 |
| **R** | 0.589 | 0.959 | 0.149 | 0.719 | 0.891 |
| **R** | 0.309 | 0.783 | 0.759 | 0.923 | 0.849 |
| **A** | 0.337 | 0.716 | 0.032 | 0.666 | 0.757 |
| **R** | 0.506 | 0.613 | 0.113 | 0.203 | 0.406 |

**Table S3. Paired (2-tail) t-tests comparing RMSFs for original $LA^-$ bound with variant $LA^-$ bound spikes' furin cleavage sites spikes**. RMSF data comparing the (protein) values for the furin site residues between different variants with $LA^-$ bound, to assess differences. Note that the lowest values indicate the greater significance for the differences between the two datasets compared. Each chain was treated separately so there were nine values for each of the residues over the 3 replicate 200 ns MD simulations for each LA-bound variant. There was weak significance between the $LA^-$bound original and LA-bound alpha for residues XR---. There was strong significance (> 95%) for differences between the RMSFs of the LA-bound original and LA-bound delta residues X-R--- and greater that 80% confidence that the RMSF for residues ---AR were also different. There is strong significance (> 95%) for the difference between residues -R-AR between the original $LA^-$ bound and $LA^-$ bound delta plus. The difference between the first, third and last furin cleavage site residues in the original $LA^-$ bound and $LA^-$ bound omicron BA.1 is strongly significant ( > 98%) and > 90% for the second and fourth residues, but the values suggest that $^{LA-}$may be de-stabilising this region. The differences between the original and omicron BA.2 furin cleavage site residues with $LA^-$ bound were only weakly significant, suggesting that the de-stabilising effect for BA.1 is not apparent in BA.2 and that $LA^-$ is having little effect on fluctuations here.

| Residue | Original-LA with Alpha-LA | Original-LA with Delta-LA | Original-LA with Delta plus-LA | Original-LA with Omicron-BA.1-LA | Original-LA with Omicron-BA.2-LA |
|---|---|---|---|---|---|
| **X** | 0.055 | 0.006 | 0.445 | 0.018 | 0.261 |
| **R** | 0.032 | 0.495 | 0.048 | 0.061 | 0.528 |
| **R** | 0.352 | 0.032 | 0.568 | 0.005 | 0.298 |
| **A** | 0.822 | 0.144 | 0.000 | 0.097 | 0.197 |
| **R** | 0.411 | 0.107 | 0.027 | 0.010 | 0.168 |
